# Population genetics of recent colonization suggests the importance of recurrent immigration on remote islands

**DOI:** 10.1101/2020.12.22.424061

**Authors:** Daisuke Aoki, Shin Matsui, Mari Esashi, Isao Nishiumi, Junco Nagata, Masaoki Takagi

**Affiliations:** Department of Natural History Science, Graduate School of Science, Hokkaido University, Sapporo, Hokkaido, Japan 060-0810; Department of Wildlife Biology, Forestry and Forest Products Research Institute, Tsukuba, Ibaraki, Japan 305-8687; Department of Biology, School of Biological Sciences, Tokai University, Sapporo, Hokkaido, Japan 005-8601; Department of Zoology, National Museum of Nature and Science, Tokyo, Japan 110-8718

**Keywords:** Bull-headed Shrike, founder effect, long-distance dispersal, island biogeography, population genetics, recurrent immigration, seasonal migration

## Abstract

**Aim:** Founder effects and recurrent immigration are two major factors that potentially contribute to genetic differentiation and population persistence in the early-stage of remote island colonization. However, their relative importance remains controversial. By conducting population genetics analyses of multiple remote island populations of the bull-headed shrike established naturally within several decades, we examined the relative contributions of founder effects and recurrent immigration on these island populations.

**Location:** Japan

**Taxon:** *Lanius bucephalus*

**Methods:** We used 15 microsatellite loci to analyze the population genetics of four newly established island populations and five Japanese mainland populations. Allelic richness, heterozygosity, genetic differentiation, and the strength of the genetic bottleneck were compared among the islands. Two analyses, STRUCTURE and the DAPC, were conducted to assess the relative influence of founder effects and recurrent immigration on genetic differentiation. Temporal samples collected over eight years on Minami-Daito Island were used to detect any change in genetic structure due to recurrent immigration.

**Results:** The founder effect strongly influenced genetic differentiation on the most remote oceanic island, Chichi-jima Island. However, this population became extinct 20 years after colonization, possibly owing to a lack of recurrent immigration. The founder effect moderately influenced a land-bridge island, Kikai-jima Island, indicating the presence of a relatively large founder population without recurrent immigration. Surprisingly, another distant oceanic island, Minami-Daito Island, was likely subject to multiple recurrent immigration events from the mainland, which obscured any genetic differentiation previously established by the founder effect.

**Main conclusion:** Underlying the island-specific population dynamics of colonization, founder effects contributed to the genetic differentiation among the three studied island populations. Importantly, however, recurrent immigration strongly affected the population persistence and subsequent evolutionary processes of remote island populations, potentially overwhelming the founder effect. We argue the importance of recurrent immigration in highly remote island colonization, which has been previously overlooked.

## Introduction

A major goal of island biogeography is to understand the processes underlying the emergence of biota on remote islands over time (Patiño et al., 2017; Warren et al., 2015). Because unique insular biodiversity consists of many island endemics that evolved on islands after colonization, understanding the processes by which the organisms arrive on an island and genetic divergence proceeds is fundamental to determining their origin (Whittaker & Fernández-Palacios, 2007). Long-distance dispersal is regarded as the major mechanism underlying the arrival of organisms on remote islands (Whittaker & Fernández-Palacios, 2007). However, the demography and population genetics associated with such arrivals have not been clarified, although three different scenarios have been proposed. The classic proposal, i.e., the founder effect scenario, suggests that a population genetic bottleneck is associated with a strong founder effect at the time of colonization, leading to rapid genetic divergence (Barton & Charlesworth, 1984). This scenario has not been supported empirically in a natural system because a severe founder effect has not been inferred in previous population genetics analyses of island colonization events. However, the two alternative scenarios explain the lack of empirical support for the classic scenario. In the second scenario, i.e., the large founder scenario, a relatively large founder population at a single colonization event results in a weak founder effect, leading to the establishment of an island population without marked genetic differentiation (Clegg et al., 2002; Pruett & Winker, 2005; Estoup & Clegg, 2003; Vincek et al., 1997). In the third scenario, i.e., the recurrent immigration scenario, recurrent immigration at the early-stage of population establishment obscures the initial founder effect, resulting in the establishment of an island population with low genetic differentiation (Grant, Grant, & Petren, 2001). These three scenarios are discriminated by (1) whether a founder effect is sufficiently influential that it results in genetic differentiation by introducing rare allele combinations and (2) whether recurrent immigration is sufficiently important that it alters the genetic pattern produced by the initial founder effect. Because founder effects and recurrent immigration have been long considered fundamental demographic processes that potentially affect the way in which founder organisms evolve (Mayr, 1954), it is important to determine which scenarios best explain colonization events.

Discriminating among various scenarios is important in island biogeography because different evolutionary processes (e.g., natural selection and genetic divergence) may follow different scenarios. For example, with a single strong founder effect, the rise of novel allelic combinations could be important for the progression of natural selection and subsequent divergence (Mayr, 1954; Barton & Charlesworth, 1984). In a large founder population, genetic drift subsequent to island colonization provides an important genetic background for adaptive evolution (Clegg et al., 2002; Sendell-Price et al., 2021). Under the strong influence of recurrent immigration, evolutionary processes that interact with gene flow, such as divergence-with-gene-flow (Feder, Flaxman, Egan, Comeault, & Nosil, 2013; Smadja & Butlin, 2011), can become important.

Distinguishing the three aforementioned scenarios has been challenging owing to the lack of a suitable system wherein a population reflecting the initial population genetic structure can be studied. Many previous studies applied genetic methods to old island colonization events, i.e., older than several hundreds of years; thus, technical difficulties in accurately estimating the timing of gene flow and genetic bottlenecks (Smadja & Butlin, 2011) have led to difficulties distinguishing the different demographic scenarios. Recurrent immigration within a short period after colonization obscures the initial founder effect (Grant et al., 2001) as the genetic consequence resembles a large founder population at a single time point (Grant, 2002). A pattern generated by a founder effect also resembles prolonged genetic drift after population establishment by a large founder population (Clegg et al., 2002). Therefore, the direct assessment of population genetic structure following recent island colonization is crucial. However, given the rarity of such colonization events, these assessments have not been achieved previously in the wild, except for a single colonization event (Grant et al., 2001) and excluding possible cases of human-assisted colonization, such as that of monarch butterflies colonized the Pacific islands (Hemstrom, Freedman, Zalucki, Ramírez, & Miller, 2022). A single case of island colonization cannot be generalized; therefore, genetic studies on multiple cases are required.

In the present study, we used a valuable study system in which the bull-headed shrike (*Lanius bucephalus*), a medium-sized passerine, has naturally colonized five remote Japanese islands and established resident populations over 50 years (Figure 1a; Table 1). These five islands are located 200–500 km beyond the normal breeding range of the species, reflecting disjunct distributions well beyond the normal dispersal range of many other medium-sized passerine species (Paradis et al., 1998), suggesting that colonization was accomplished via rare long-distance dispersal events. Two islands, namely Nakano-shima Island and Kikai-jima Island, (KKJ) are land-bridge islands in the Ryukyu Archipelago (Japan’s southwestern most islands) where shrikes are rare seasonal migrants (Amami Ornithologists’ Club, 1997; Okinawa Wild Bird Study Group, 1986). In contrast, Minami-Daito Island (MDT), Kita-Daito Island (KDT), and Chichi-jima Island (CCJ) are oceanic islands on the Pacific Sea, where rare vagrant birds can arrive during spring and autumn (Takehara, Anezaki, Takagi, Okudo & Knagawa, 1999; Ando, Emura & Deguchi, 2020). Therefore, population genetics analyses of these multiple remote islands will potentially reveal variation in colonization scenarios with early-stage population genetic structures. Scenario comparisons of multiple islands will likely lead to both generalizable and island-specific results, allowing us to evaluate the influence of founder effects and recurrent immigration. The uniqueness of our study system is further improved by different colonization consequences on two of the oceanic islands: the CCJ population became extinct 20 years after population establishment, whereas the MDT population has persisted for roughly 50 years. Therefore, we are able to infer a relationship between the influence of the founder effect and recurrent immigration and the fate of the established populations.

**Figure 1.**
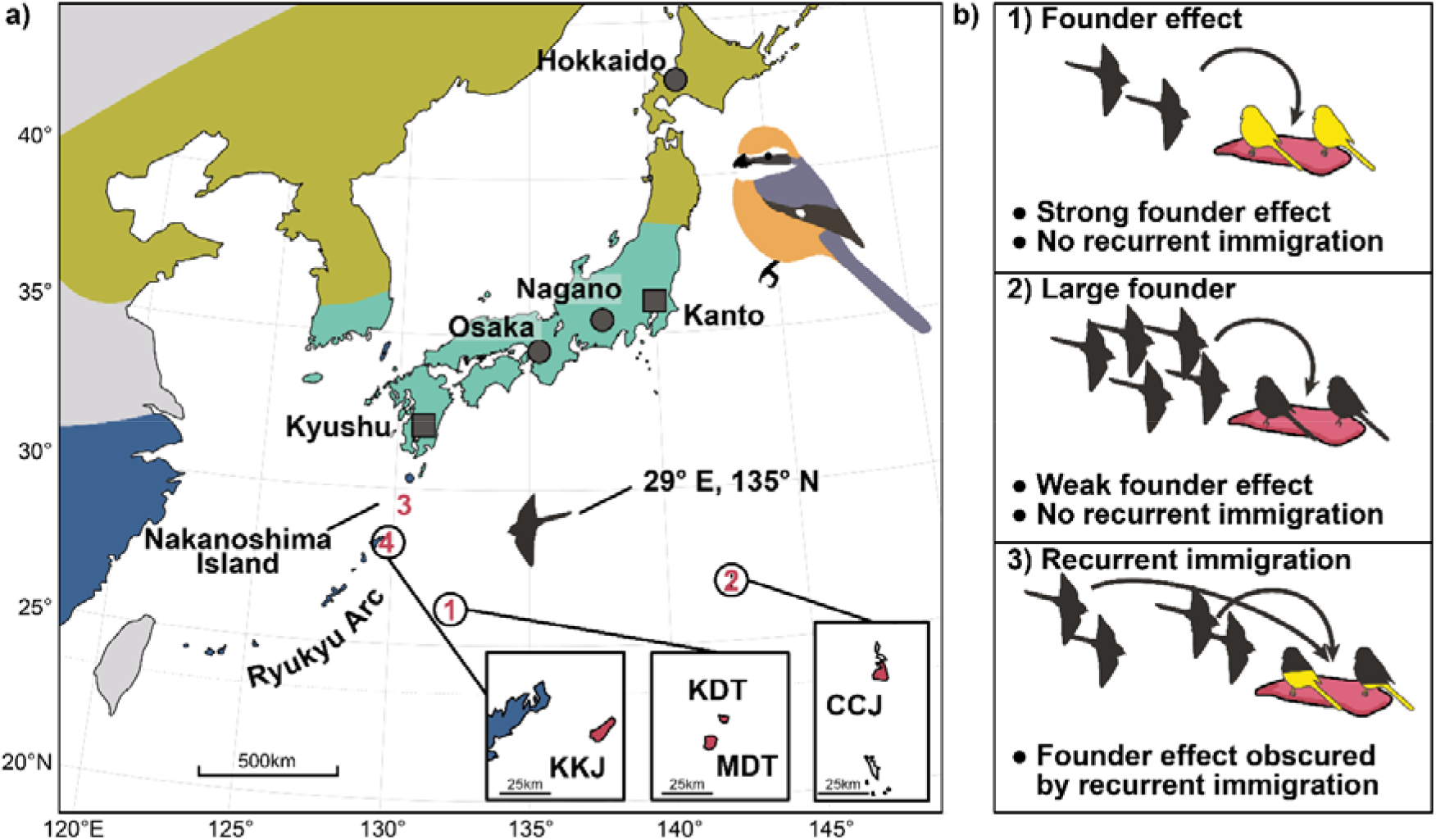
(a) The range of the bull-headed shrike and the location of sampling sites for this study. Land colours correspond to different origins of the populations: summer migrants (yellow), both residents and migrants (green), winter visitors or rare seasonal migrants (blue), and residents due to recent colonisation (red, the numbers correspond to the colonisation sequence). Sampling sites are denoted by circles (intensive local sampling) and squares (opportunistic sampling). (b) The three different scenarios with different contributions by the founder effect and recurrent immigration are shown. Abbreviations for island populations are as follows: Chichi-jima = CCJ, Kikai-jima = KKJ, Minami-Daito = MDT, and Kita-Daito = KDT.

**Table 1.**
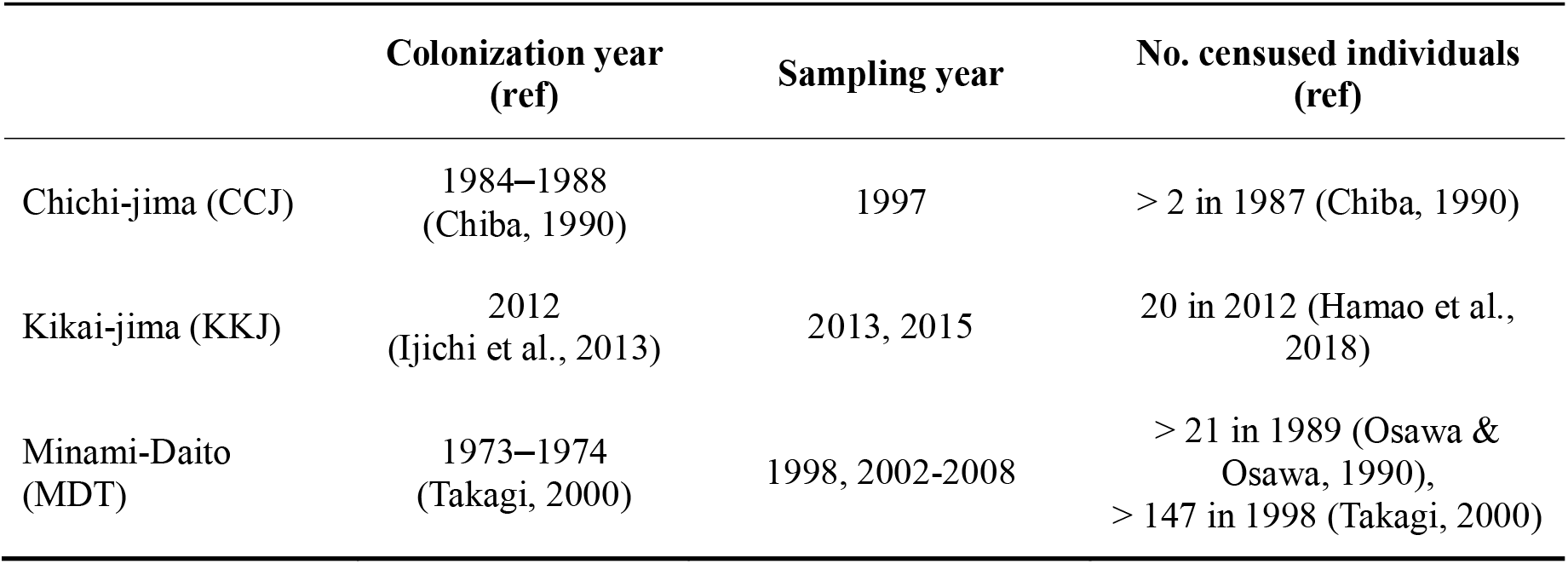
Summary for the details of island colonization and sampling years on the three different islands

|  | Colonization year<br>(ref) | Sampling year | No. censused individuals<br>(ref) |
| --- | --- | --- | --- |
| Chichi-jima (CCJ) | 1984–1988<br>(Chiba, 1990) | 1997 | > 2 in 1987 (Chiba, 1990) |
| Kikai-jima (KKJ) | 2012<br>(Ijichi et al., 2013) | 2013, 2015 | 20 in 2012 (Hamao et al.,<br>2018) |
| Minami-Daito<br>(MDT) | 1973–1974<br>(Takagi, 2000) | 1998, 2002-2008 | > 21 in 1989 (Osawa &<br>Osawa, 1990),<br>> 147 in 1998 (Takagi, 2000) |

We performed population genetics analyses of bull-headed shrikes on the main islands of the Japanese Archipelago (hereafter, “the mainland”) and four remote islands using 15 microsatellites. Genetic diversity and the level of genetic differentiation were estimated using common statistical measures, e.g., heterozygosity, allelic richness, and Fst, as well as bottleneck tests, and genetic differentiation was also evaluated using Bayesian clustering analysis and multivariate analysis. We did not apply immigrant detection methods, e.g., BayesAss (Wilson & Rannala, 2003), or gene flow estimates via isolation-with-migration models, e.g., IMa (Hey, 2010), because the islands were colonized within a few to several decades, suggesting that the island populations would not satisfy the requirements of these methods, i.e., sufficient genetic differentiation (Wilson & Rannala, 2003) and/or divergence (Hey, 2010) from the populations under comparison. Nevertheless, the use of standard genetic methods to analyze the newly founded populations allowed us to make predictions under the three aforementioned scenarios. The founder effect scenario (scenario 1: Figure 1b-1) assumes a strong founder effect and no recurrent immigration; thus, an insular population representing a rare allele combination of mainland individuals is expected (Slatkin, 2004). Hence, we predicted a strong genetic bottleneck, high genetic differentiation, and low allelic richness and heterozygosity under scenario 1. The large founder scenario (scenario 2: Figure 1b-2) assumes a weak founder effect and no recurrent immigration; thus, under this scenario, we predicted no apparent genetic differentiation and no representation of the mainland allele combination, a moderate genetic bottleneck, and moderately low allelic richness because “a large founder” would not be as large as the mainland population and allelic diversity is sensitive to changes in population size (Nei, Maruyama, & Chakraborty, 1975). Reduction in expected heterozygosity is also predicted under the large founder scenario because expected heterozygosity can be restored slowly without recurrent immigration (Keller et al., 2001; Nei et al., 1975). The recurrent immigration scenario (scenario 3: Figure 1b-3) assumes that recurrent immigration obscures the founder effect; therefore, we predicted both high allelic richness and heterozygosity as well as a low level of differentiation (Grant et al., 2001; Keller et al., 2001). Temporal samples collected across eight years from MDT were available, allowing us to identify the recurrent immigration scenario, i.e., the level of genetic differentiation of the island population relative to that of the mainland should decrease after a recurrent immigration event. Based on our findings, we discuss the contribution of the founder effect and recurrent immigration to the genetic differentiation and population persistence of remote island populations following colonization.

## Methods

### Study system and field procedure

The first observations of breeding attempts by bull-headed shrikes on various remote Japanese islands are as follows: (1) on MDT and KDT, in the Daito Islands, in 1973–1974 (Takagi, 2000); (2) on CCJ, in the Ogasawara Islands, in 1984–1988 (Chiba, 1990); (3) on Nakano-shima, in the Tokara Islands, in 1989 (Morioka, 1990); and (4) on KKJ, in the Amami Islands, in 2012 (Ijichi, Torikai, & Hamao, 2013) (Figure 1; Table 1). On the Japanese mainland, shrikes are partial migrants and both migratory and resident shrikes co-occur; in contrast, in the northern part (e.g., Hokkaido) or high mountain ranges (e.g., the Japanese Alps) of Japan, a high proportion of seasonal migrants occur (Imanishi, 2005; Ministry of the Environment of Japan, 2020).

Blood samples were collected from shrikes from several populations on mainland Japan and from four of the five remote islands (MDT, KDT, CCJ, and KKJ) (Figure 1; Table 1). Samples were stored in 99.5% ethanol. Field sampling was conducted locally in a time-intensive manner for the insular populations and many of the mainland populations during the breeding period (n = 54 on Osaka in 1989; n = 32 on Hokkaido in 1998; n = 15 on CCJ in 1997; n = 4 on KDT in 2008; n = 5 on KKJ in 2013 and 2015; n = 25 on Nagano in 2019; see below for MDT). For CCJ, additional samples were collected in 1995 (n = 1) and 1998 (n = 2), and these were also included in two analyses: STRUCTURE and the discriminant analysis of principal components (DAPC). Mist nets and spring net traps were used to capture shrikes for blood sampling. Tissue and blood samples from the Kyushu (n = 14) and Kanto (n = 18) regions were collected opportunistically, resulting in wider regional sampling over multiple years. Samples in these two regions were collected both in the breeding seasons and prebreeding seasons when shrikes occupy breeding territories for the coming spring (Kurata, 1967). Although opportunistic sampling can affect genetic results, the effect of different sampling schemes on our results was limited. On MDT, samples from multiple breeding seasons, i.e., in 1998 (n = 30) and annually from 2002 to 2008 (n = 365 in total), were collected. We used the samples collected in 1998 as representative of the earlier genetic structure of the MDT population for most of the genetic analyses, whereas data collected in 2002–2008 were combined with those from 1998 to analyze the temporal genetic change. See Table S1 for sample details.

### Laboratory procedure and calculation of genetic diversity indices

DNA was extracted from samples using either a Qiagen Blood & Tissue Kit (Qiagen) or Dr.GenTLE (TaKaRa) following the manufacturers’ protocols. The genotype of each of the samples was determined at 15 microsatellite loci. We followed the method of Matsuo et al. (2014) in our experimental protocols. The primers used in the present study are summarized in Table S2. Because each population could include a different proportion of closely related individuals, the results of the following genetic analyses may be biased (Devlin & Roeder, 1999). Therefore, we modified a dataset in which the relatedness of individuals was controlled for each population by retaining only one individual for each full-sib cluster inferred using COLONY v.2.0.6.6 (Jones & Wang, 2010). We used the dataset without relatedness in the following analyses. See the supplementary methods for full details.

Null allele frequencies were estimated for each locus for each population using FreeNA (Chapuis & Estoup, 2007). Tests for linkage disequilibrium across all the populations were conducted using GENEPOP v. 4.7.5 (Raymond & Rousset, 1995; Rousset, 2008). We compared allelic richness, expected heterozygosity, and observed heterozygosity among populations by constructing linear mixed regression models using the ‘lmerTest’ package in R (Kuznetsova, Brockhoff, & Christensen, 2017). We separately constructed a model for each set containing one island (CCJ, KKJ, or MDT) and five mainland regions to estimate the degree of reduction in the genetic diversity indices (KDT was excluded because it was considered a sink population of the MDT population based on the DAPC). In each model, we assigned the five mainland regions as “mainland” and one island as “island” and set these as the explanatory variables, whereas the locus identity was set as a random factor. We estimated a model coefficient of the effect of the category “island” and its statistical significance in each model. Allelic richness was calculated according to the rarefaction method using HP rare (Kalinowski, 2004, 2005), which performs rarefaction for unbiased estimates of allelic richness. Given that the smallest sample size was four (KDT), it was rarefied to eight genes per locus. We calculated expected and observed heterozygosity for each locus for each population under two different models: model one accounted for the presence of null alleles and genotyping failures, whereas model two also accounted for the inbreeding coefficient of the first model determined using INEST v. 2.2 (Chybicki & Burczyk, 2009). The model with the lowest deviance information criterion (DIC) value may outperform the other. Chains with 1,000,000 cycles and a burn-in of 100,000 cycles were run, and parameters were retained every 100th update. As the inbreeding coefficient was not significant in any population (see Results), heterozygosity calculated under the first model was compared among populations.

A test for heterozygosity excess under the two-phase model (Cornuet & Luikart, 1996) and the *M*-ratio test (Garza & Williamson, 2001) were conducted as genetic bottleneck tests for each population in INEST v. 2.2. An *M*-ratio of <0.68 was determined as the signature of a bottleneck effect (Garza & Williamson, 2001). Pairwise Fst values between pairs of populations were calculated with 95% confidence intervals, and the presence of null alleles was accounted for using FreeNA (Chapuis & Estoup, 2007). Differences in mean Fst values were compared between each set of inter-mainland–island comparisons for each island (e.g., between the Fst of Kanto–MDT and that of Hokkaido–MDT). Significance levels were calculated using two-sided permutation tests with 10,000 resampling iterations.

### Spatial genetic structure

STRUCTURE analysis (Falush et al., 2003; Pritchard et al., 2000) was conducted based on the “Admixture model” assuming correlated allele frequencies among populations with 10 replicates of 100,000 cycles of burn-in and 500,000 cycles of the Markov chain Monte Carlo. The number of genetic clusters, *K,* was tested from 1 to 10. The Evanno method was used to infer the best value of *K* according to Δ*K* values from the results (Evanno, Regnaut, & Goudet, 2005). Ten replicates were combined into one output using CLUMPP (Jakobsson & Rosenberg, 2007), and the results are shown across several *K* values including the best *K*. The DAPC (Jombart et al., 2010) was implemented via the R package ‘adegenet’ v. 1.2.1 (Jombart, 2008) to assess genetic structure within a complex population structure, which was suspected in the studied populations based on STRUCTURE analysis. We performed 20,000 replicates of cross-validation to determine the number of principal components with the lowest mean squared error to be retained in the DAPC. After cross-validation, 32 principal components were retained in the DAPC, and the first and the second discriminant functions (DA1 and DA2) were used for plotting.

### Temporal samples for MDT

We reperformed the STRUCTURE analysis and the DAPC with additional samples collected in 2002–2008 on MDT to make further predictions. For the STRUCTURE analysis, we used the option “PFROMPOPFLAGONLY” with *K* = 4. The DAPC also has an option to predict additional samples using the function “predict.dapc.” Parameter settings for these analyses are described in the supplementary methods. The temporal change in the genetic structure on MDT was assessed by conducting Mantel tests on samples collected in different years using ‘ape’ v. 5.3 (Paradis & Schliep, 2019) based on pairwise Fst and Cavalli-Sforza and Edwards’ genetic distance Dc (Cavalli-Sforza & Edwards, 1967) calculated for samples collected in different years using FreeNA.

## Results

### Genetic diversity indices

In total, 565 individuals were genotyped for 15 microsatellite loci; only 0.76% of the dataset was missing data. After conducting sib-ship assignment analysis, 85 related individuals were removed, and the dataset included 177 individuals from five mainland and four island populations (the representatives of MDT were those individuals sampled in 1998). After conducting the Bonferroni correction for multiple comparisons, there was no evidence of linkage equilibrium. There were no alleles for which the null allele frequencies were >0.2 across all populations (Table S3). In the mixed linear model comparisons between the mainland populations and one island population, allelic richness was significantly lower for all insular populations, although the degree of such a reduction was lowest for MDT, moderate for KKJ, and highest for CCJ (Figures 2a and 3a). A pattern was similar for observed heterozygosity, although the observed reduction was not statistically significant for the MDT and KKJ populations (Figure 3c). Contrastingly, the reduction in expected heterozygosity was significant for the CCJ and KKJ populations but not the MDT population (Figures 2b and 3b).

**Figure 2.**
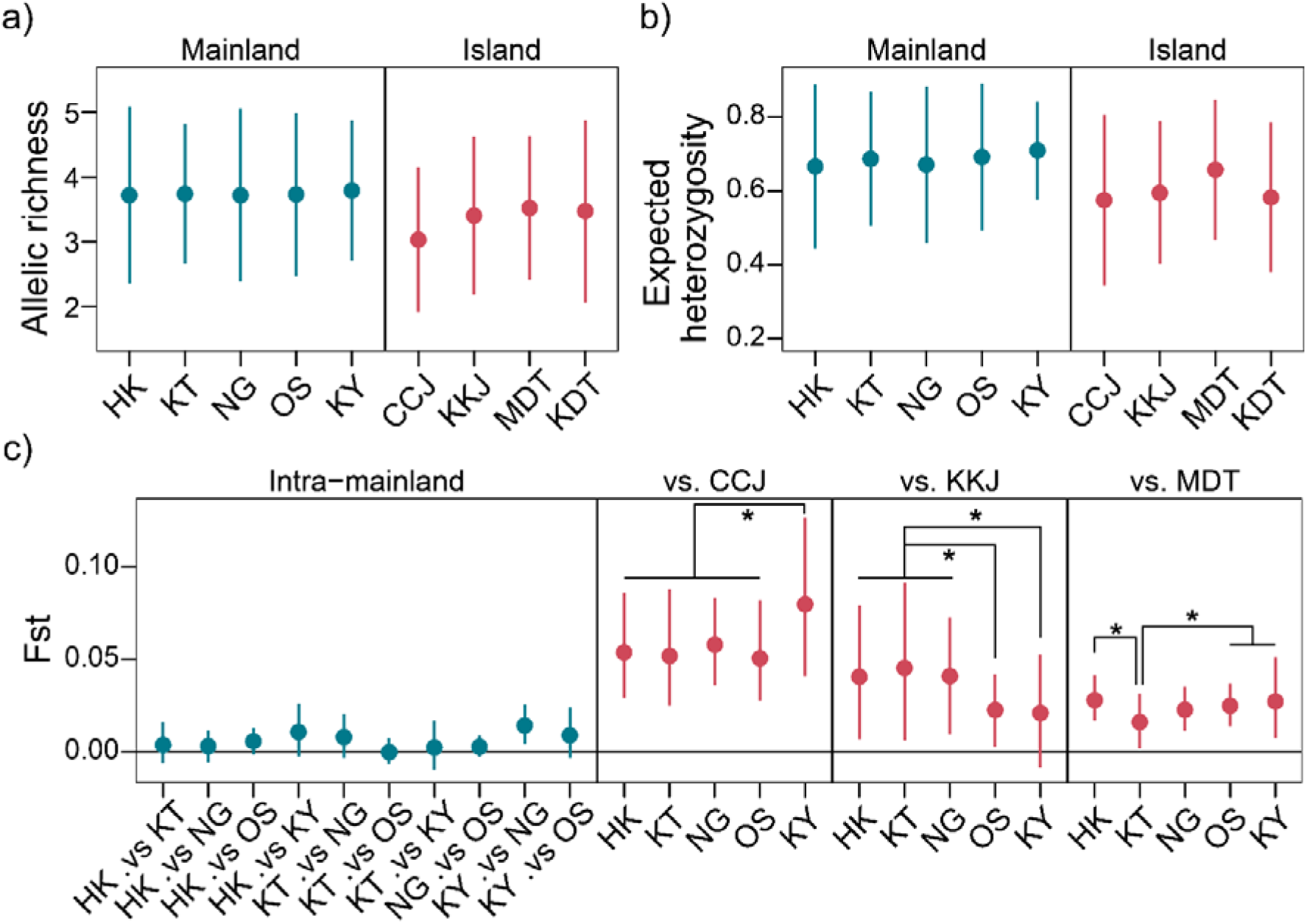
Comparisons of genetic diversity indices across five mainland and four insular populations, including (a) allelic richness corrected by rarefaction, (b) expected heterozygosity, and (c) pairwise Fst. Mean values with 95% confidence intervals across 15 loci were indicated. Results of permutation tests were indicated for pairwise Fst within each category of island-mainland comparisons (comparisons with *p* < 0.005 were indicated with asterisks). Abbreviations for populations are as follows: Hokkaido = HK, Kanto = KT, Nagano = NG, Osaka = OS, Kyushu = KY, and others refer to Figure 1.

**Figure 3.**
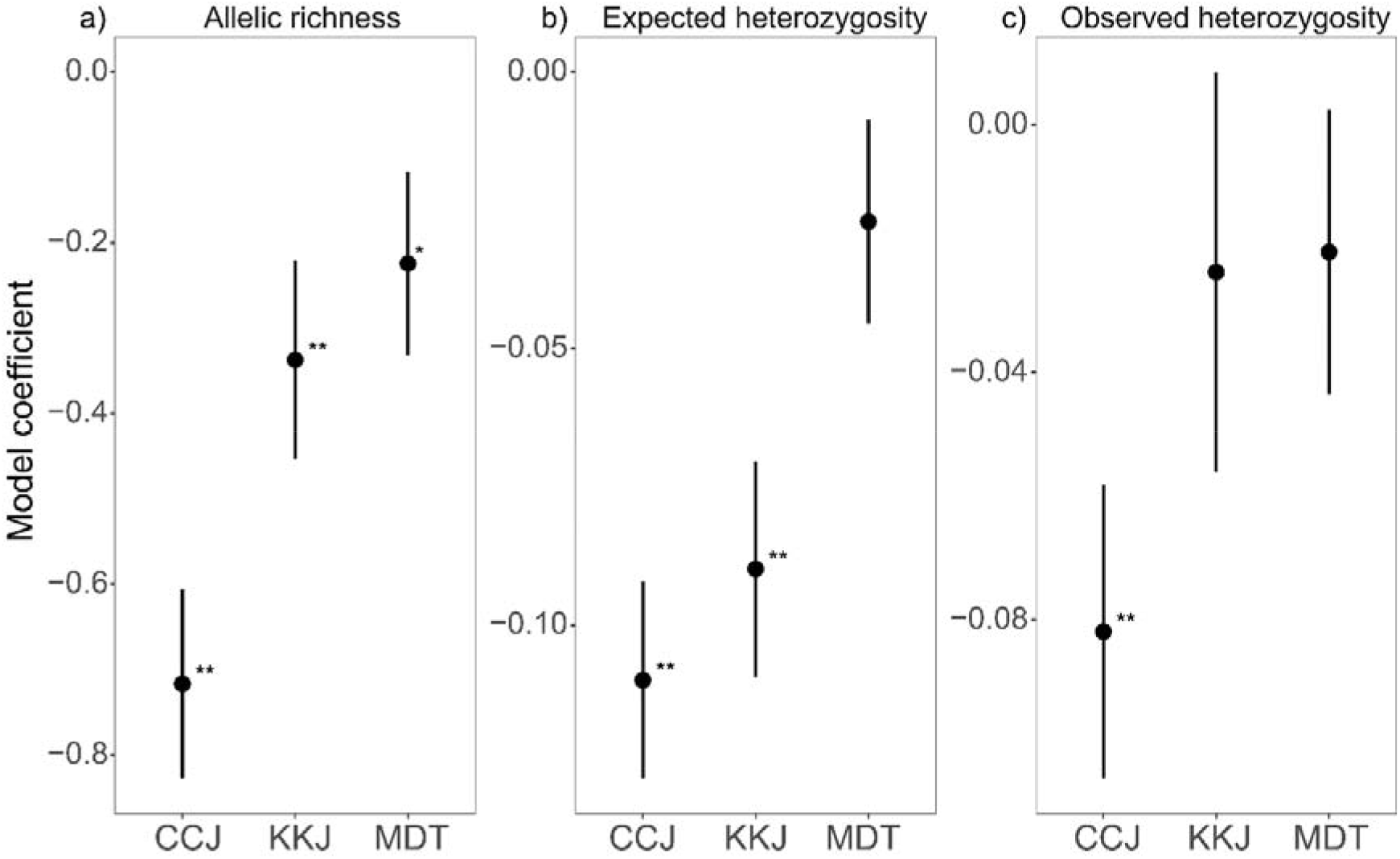
Estimated model coefficients of the effects of island on the three different genetic diversity indices from the linear mixed models constructed for each island populations. Coefficients indicate the level of reduction in the indices when compared to the mainland populations, and asterisks denote their statistical significance (* 0.01 ≤ *p* < 0.05, ** *p* < 0.01).

A model including the inbreeding coefficient performed better than a model without this coefficient only in the Kyushu population based on DIC values (Table 2), whereas the 95% credibility intervals of inbreeding coefficients included zero for all populations, indicating that inbreeding was not supported statistically in any population. A population bottleneck was supported statistically for the CCJ and KDT populations by both the heterozygosity excess and *M*-ratio test (Table 2). The *M*-ratio was <0.68 for the KKJ population; given that the *M*-ratio is more sensitive for detecting a genetic bottleneck than testing a heterozygosity excess (Cornuet & Luikart, 1996; Garza & Williamson, 2001), a genetic bottleneck in the KKJ population was supported to a lesser extent than that in the CCJ population.

**Table 2.** Tests for inbreeding and genetic bottlenecks in mainland and insular bull-headed shrike populations. The sample size (*N*) used for calculation is provided for each population. A population for which a model considering inbreeding performed better than one without it is indicated with “+”. Inbreeding coefficient was calculated under a model with inbreeding. For the statistics for genetic bottlenecks, *p*-values calculated by Wilcoxon signed-rank test based on 1,000,000 permutations for heterozygosity (Hz) excess and *M*-ratio are given. All the indices were calculated by INEST v. 2.2 (Chybicki and Burczyk, 2009)

| | $N$ | Inbreeding Model | Inbreeding Coefficient [95% credibility interval] | $P$ (Hz excess) | $M$-ratio |
| --- | --- | --- | --- | --- | --- |
| **Mainland** |  |  |  |  |  |
| Hokkaido | 32 | - | 0.018 [0, 0.05] | 0.92 | 0.99 |
| Kanto | 18 | - | 0.031 [0, 0.08] | 0.99 | 0.99 |
| Nagano | 21 | - | 0.031 [0, 0.09] | 0.78 | 0.86 |
| Osaka | 42 | - | 0.029 [0, 0.07] | 0.96 | 1.0 |
| Kyushu | 14 | + | 0.063 [0, 0.12] | 0.90 | 0.86 |
| **Island** |  |  |  |  |  |
| Chichi-jima (CCJ) | 15 | - | 0.026 [0, 0.08] | **0.014\*** | **0.003**<sup>□</sup> |
| Kikai-jima (KKJ) | 5 | - | 0.027 [0, 0.1] | 0.13 | **0.48**<sup>□</sup> |
| Minami-Daito (MDT) | 23 | - | 0.023 [0, 0.07] | 0.73 | 0.81 |
| Kita-Daito (KDT) | 4 | - | 0.025 [0, 0.1] | **0.014\*** | **0.43**<sup>□</sup> |

\* $p < 0.05$ , <sup>□</sup> $M$ -ratio $< 0.68$

### Genetic differentiation and spatial genetic structure

The genetic differentiation of insular populations was supported by Fst values, STRUCTURE analysis, and the DAPC, although the degree of genetic differentiation differed among populations (Figure 2c; Figure 4a, b). The differentiation of insular populations from the mainland populations was statistically supported by the Fst values of most pairs. The degree of genetic differentiation of the island populations relative to the mainland populations was higher for CCJ than for KKJ and MDT (*p* = 0.0078 and 0.01 for KKJ and MDT, respectively; 10,000 permutations; α = 0.017 after Bonferroni correction). The degree of differentiation of each island population also varied depending on which mainland populations were used for comparison; for instance, a lack of genetic differentiation between the KKJ and Kyushu populations and low differentiation between the MDT and Kanto populations were inferred (10,000 two-tailed bootstrap; α = 0.005 after Bonferroni correction; Figure 2c).

**Figure 4.**
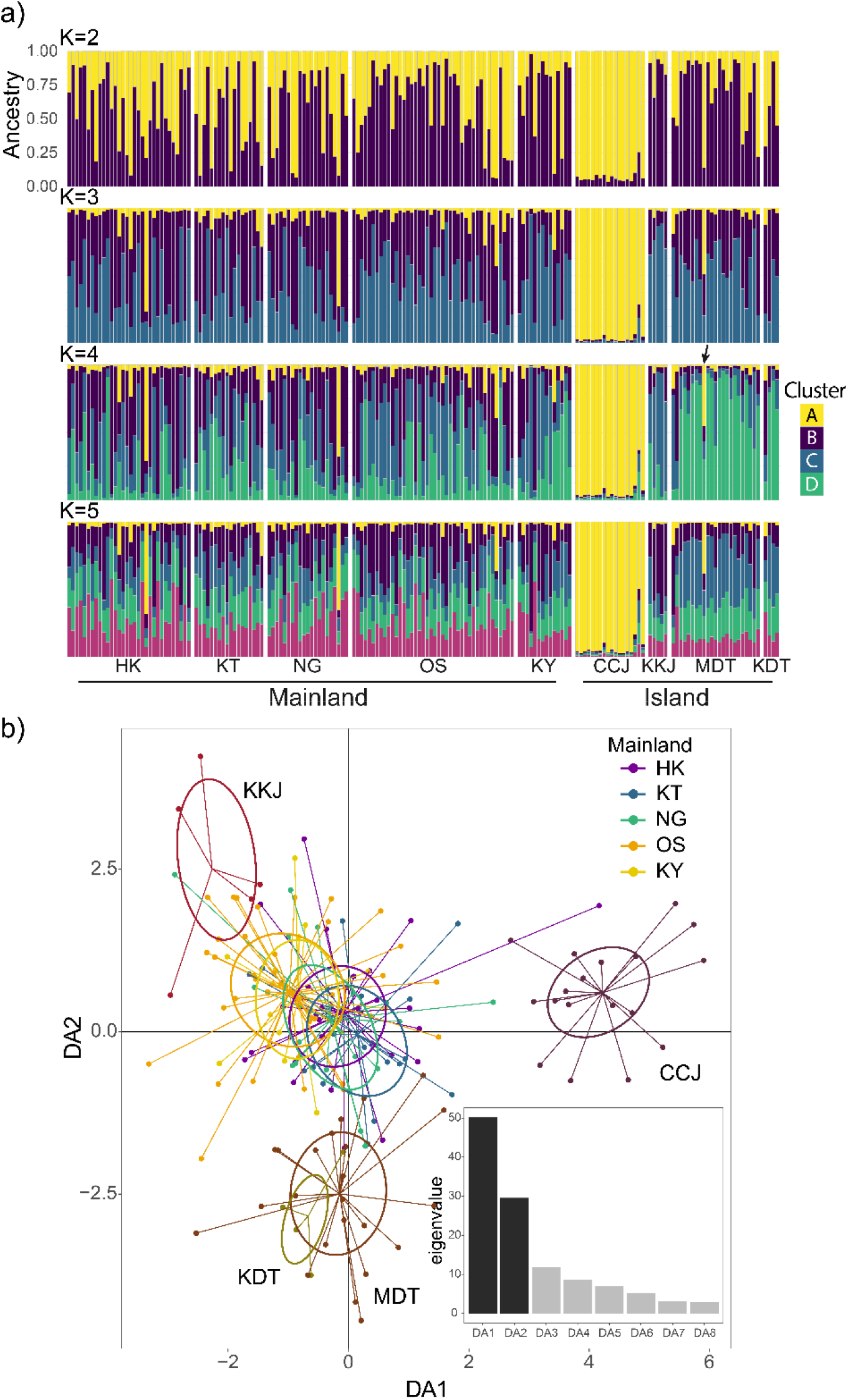
Results of genetic clustering for the mainland and insular individuals, inferred by the (a) STRUCTURE and (b) discriminant analysis of principal components (DAPC). (a) Genetic assignment of individuals to each number of genetic cluster *K,* ranging from *K* = 2 to 5, is shown (the highest Δ*K* occurred at *K* = 4). Each bar indicates an individual, and the height of different colours shows the assignment probabilities to each corresponding genetic cluster. (b) A scatterplot shows the first two discriminant functions of the DAPC. Different colours of dots and inertia ellipses represent different sampled populations. In an inset, eigenvalues are shown for each discriminant function.

The apparent genetic differentiation of the CCJ population was supported by STRUCTURE analysis (with the highest *ΔK* values occurring at *K* = 4; Figure S1) (Figure 4a) and the DAPC (the first discriminant function with the highest discriminatory power; Figure 4b). At *K* = 4 in STRUCTURE analysis, most individuals were assigned to a major genetic cluster in CCJ with probabilities of >80% (cluster A [yellow] in Figure 4a). Although cluster A was rare in all mainland populations, a few individuals were associated with this genetic cluster with higher probabilities, i.e., one from Hokkaido (68.3%) and one from Nagano (65.3%). No individual from Kanto, Osaka, or Kyushu exceeded a 50% assignment probability to cluster A. The DAPC gave similar results, i.e., the position of one Hokkaido sample assigned largely to cluster A was inferred with a first discriminant score as large as that of the CCJ population. Therefore, it is likely that only a small number of shrikes inhabiting Hokkaido (or a population nearby) colonized CCJ, leading to a strong founder effect and fixation; thus, the CCJ population is genetically differentiated and reflects the rare allelic combination of the mainland.

In 1998, the MDT population exhibited a skewed genetic cluster composition and slight genetic differentiation according to STRUCTURE analysis (cluster D [green] was the major cluster) and could also be discriminated via the second discriminant function of the DAPC (Figure 4). Genetic clusters were shared with many mainland individuals in STRUCTURE analysis, and more individuals from MDT were located toward the mainland population cluster than those from CCJ and KKJ in the DAPC plane. Furthermore, STRUCTURE analyses including temporal samples collected between 1998 and 2008 indicated that an increase in cluster C (blue) and decrease in cluster D (green) occurred between 1998 and 2002 (Figure 5a), corresponding to the shift toward positive DA2 scores in the DAPC (Figures 5b and S2), indicating a replacement of the major genetic cluster with that from the mainland. Results of the Mantel test supported a temporal change of genetic structure on MDT over the eight-year sampling period based on both the Fst (*R* = 0.70, *p* = 0.001) and Cavalli-Sforza and Edwards’ genetic distance (*R* = 0.60, *p* = 0.032) (Figure S3), whereas the pattern was weakened (Fst: *R =* 0.40, *p* = 0.037) or statistically unsupported (Cavalli-Sforza and Edwards’ genetic distance: *R* = 0.15, *p* = 0.25) when the samples from 1998 were excluded (Figure S4). No change in heterozygosity or allelic richness was observed throughout the study years on MDT (Figure S5). Together, these results indicate that weak genetic differentiation was once established on MDT but was obscured between 1998 and 2002 by recurrent immigration and gene flow from the mainland. Notably, only one individual had cluster A at a relatively high percentage (38.5%; Figure 4a, arrow). The KDT population was located near the MDT population on the DAPC plane, suggesting that a metapopulation structure exists between these two islands in the Daito Islands, i.e., that KDT is the sink population. This interpretation is corroborated by strong support for the bottleneck (Table 2).

**Figure 5.**
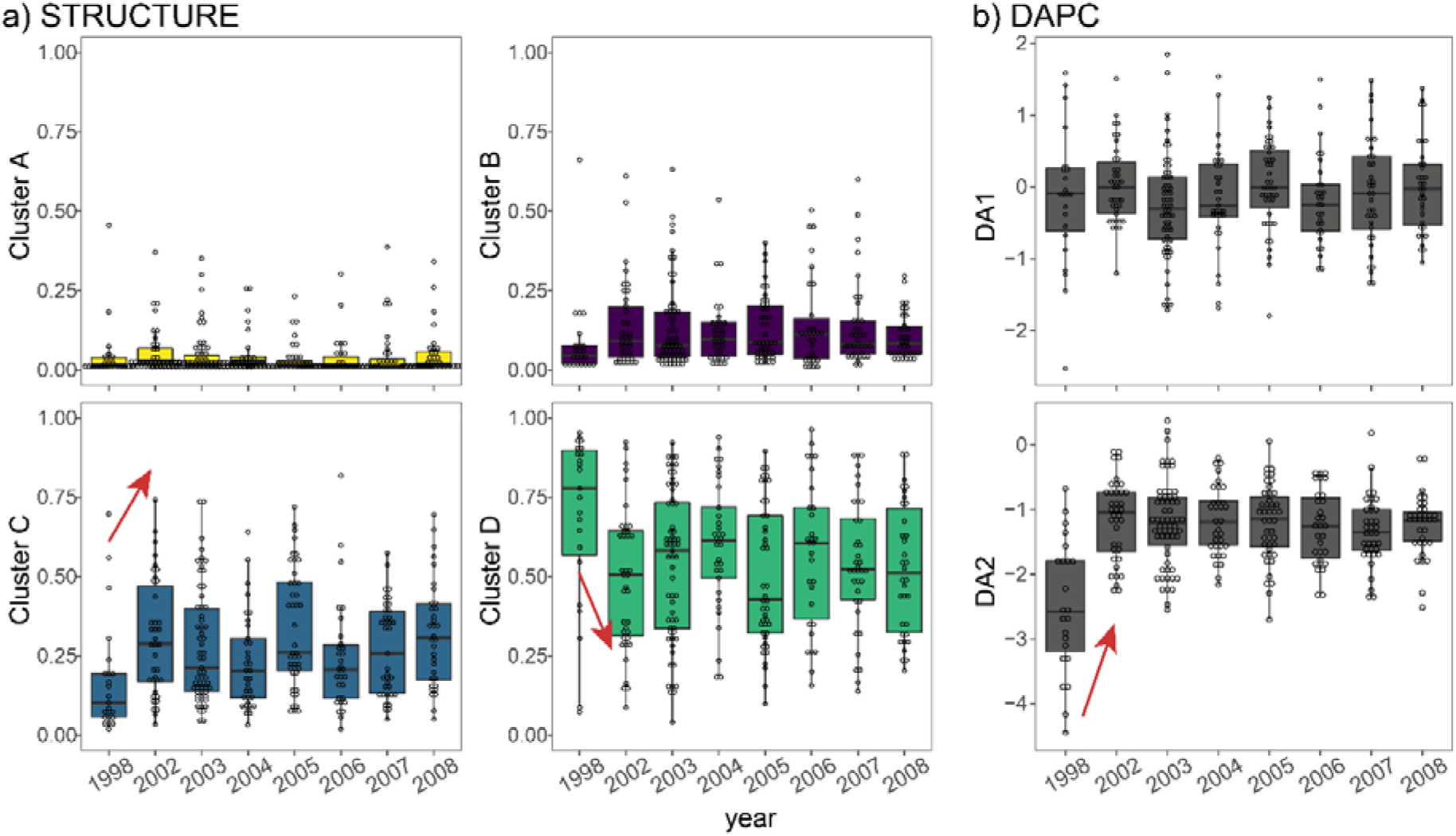
Temporal changes in the genetic compositions of the Minami-Daito (MDT) island population. The y-axes represent either (a) the percentages of each genetic cluster in the STRUCTURE analysis and (b) the positions at each discriminant functions in the DAPC. Each dot represents individuals collected in the corresponding years, and their population-level median and interquartile ranges are indicated. Note a temporal change between 1998 and 2002 (indicated by red arrows).

In the STRUCTURE analysis, the genetic distinctiveness of the KKJ population was inconclusive. Based on the discriminant functions of the DAPC, there was a slight trend indicating that eastern, northern, or high-altitudinal populations (Hokkaido, Kanto, and Nagano) were located toward the fourth quadrant, whereas western populations (Osaka and Kyushu) were located toward the second quadrant of the plane. Therefore, the genetic differentiation of the KKJ population was less pronounced than that of the western mainland populations.

## Discussion

In the present study, population genetics analyses of multiple island populations of the bull-headed shrike supported different scenarios of colonization for different islands. Analysis results of the CCJ population were concordant with the founder effect scenario (scenario 1), i.e., a genetic structure representing a rare genetic cluster on the mainland, strong signatures of a genetic bottleneck, and significantly lower allelic richness and heterozygosity than those of the mainland. Despite the small sample size of the KKJ population, its allelic richness was significantly lower than that of the mainland yet higher than that of the CCJ population, and the genetic bottleneck signature was weaker than that of the CCJ population. The genetic differentiation was slight from most of the mainland populations and was not even supported from the possible source area (Kyushu), indicating that scenario 1 was unlikely (Clegg et al., 2002; Estoup and Clegg, 2003). In addition, significantly low expected heterozygosity i.e., low allele frequency, allowed us to reject the recurrent immigration scenario (scenario 3). Therefore, the large founder scenario (scenario 2) was most likely for the KKJ population. The inferred difference in the influence of founder effects on the CCJ and KKJ populations is unlikely to be an artifact of our sampling scheme because (1) a significant difference in the initial population sizes of the two islands was found in field censuses (Table 1) and (2) the mean expected heterozygosity of the KKJ population was similar to that of *Zosterops lateralis* populations without severe bottlenecks, whereas that of the CCJ population was as low as that of *Zosterops* populations with strong bottlenecks (Clegg et al., 2002). In contrast, analysis results of the MDT population in 1998 were concordant with scenario 3, i.e., supported by the lack of a genetic bottleneck, a heterozygosity as high as that of the mainland populations, relatively high allelic richness, and a low level of genetic differentiation. Recurrent immigration also possibly occurred between 1998 and 2002, further obscuring genetic differentiation. These findings support the occurrence of at least two recurrent immigration events since population founding and before 2008. Overall, our genetic analyses of multiple new island populations suggest that island populations can be established through different combinations of the founder effect and recurrent immigration.

Although differences between the recurrent immigration and large founder scenarios have been debated (Clegg et al., 2002; Grant, 2002), they may not be mutually exclusive. The abrupt change in genetic structure on MDT between 1998 and 2002 suggests that recurrent immigration occurred in a large flock. This notion is supported by the contrary case of cluster A, which did not spread to MDT after 1998 while only one breeding female shrike was found with this cluster at 38% of its genetic contribution in 1998. Moreover, the low but statistically significant genetic differentiation of the MDT population in 1998 could only be established through a founder effect. Collectively, these results indicate that the MDT population may have been established by a relatively large founding population, which did not result in the representation of a rare allele combination but was sufficient to skew the genetic composition, which was in turn obscured by subsequent multiple recurrent immigrations. Multiple immigration events highlight the scenario wherein a founder population does not necessarily consist of one discrete and simultaneous arrival but rather several arrivals that continue over time. This scenario is supported by a direct observation of colonization by Galápagos finches (Grant et al., 2001), wherein breeding was initiated after years of continuous visits by immigrants upon colonization. The case on the Galápagos Islands involved colonization from source populations several tens of kilometers away; however, this scenario may be prevalent even under geographically remote conditions, like in our case. Thus, a founder population, which usually assumes temporal discreteness from recurrent immigrants (Mayr, 1954), may be difficult to define owing to temporal continuity (Grant, 2002). Nevertheless, temporal dynamics in the number of immigrants may exist, and arrival peaks may be concentrated within a few years, as shown in our temporal MDT analyses and by Grant et al. (2001). Therefore, despite difficulties in distinguishing “founder individuals” and “recurrent immigrants” at the individual level, we successfully evaluated their genetic influence at the temporally concentrated “immigration cluster” level. From this perspective, the newest island population on KKJ may not have reached the timestep at which it receives a recurrent immigrant cluster.

The absence of recurrent immigration is also thought to have influenced the population persistence of the CCJ population. In our analysis results for CCJ, strong genetic differentiation driven by sampling of a few mainland individuals indicated the efficacy of the founder effect in generating genetic differentiation. However, the reduction of genetic variability in the CCJ population indicated that extinction was inevitable for this population (Lynch, Conery, & Burger 1995). Although we did not directly assess the effect of inbreeding depression on population persistence in the present study, we did identify possible inbreeding-related morphological defects in the CCJ population (D. Aoki & M. Takagi, in prep.). Moreover, the shrikes on CCJ were highly sensitive to human disturbance (M. Takagi pers. obs.), which has also been reported on KKJ and MDT (Hamao, Torikai, Yoshikawa, Yamamoto, & Ijichi, 2021). Therefore, the CCJ population was probably susceptible to genetic deficiencies and demographic and/or environmental stochasticity (or a combination of these factors), which has been inferred in cases of species invasion (Lockwood, Conery, & Blackburn, 2005). If recurrent immigration occurred before extinction, it could have resulted in the rescue effect for the CCJ population, allowing it to persist (Brown & Kodric-Brown, 1977). We suggest that a population founded once is likely to receive subsequent immigration if it persists until recurrent immigration occurs (as we have discussed for the MDT population), which is possible because island colonization is not geographically random due to the directionality of dispersal vectors such as the wind and sea current (Gillespie et al., 2012).

Together, we propose that, even in remote island colonization, the genetic impact of recurrent immigration is crucial for establishing the early-stage population genetic structure as it obscures and overcomes the initial founder effect. Previously, the contribution of recurrent immigration has been overlooked at the initial stage of population establishment because full allopatry was assumed soon after island colonization in terms of the geographic mode of speciation (Warren et al., 2015), although its demographic processes were not often considered (Harvey, Singhal, & Rabosky, 2019). Genomic studies on colonization in the evolutionary past have inferred the importance of postdivergence gene flow (e.g., Lamichhaney et al., 2015; Sendell-Price et al., 2020), although the timing of gene flow has remained controversial owing to the technical difficulty of determining this timing (Smadja & Butlin, 2011). Studies on recent colonization events involving Galápagos finches (Grant, 2002; Grant et al., 2001) and song sparrows on Mandarte Island (Keller et al., 2001) have suggested the importance of recurrent immigration for population persistence, although these populations were geographically close to the surrounding islands (several to tens of kilometers, i.e., a parapatric situation), so recurrent immigration was expected under the metapopulation framework (Hanski, 1998). In contrast, our cases included geographically remote islands (several hundreds of kilometers from the mainland source), i.e., long-distance dispersal outside the normal dispersal range. Therefore, the genetic comparisons of multiple populations in the present study provided a new and important insight into the origin of remote island populations that are allopatric in terms of speciation. Recurrent immigration counteracts founder effects to allow population persistence (Brown & Kodric-Brown, 1977), which creates the opportunity for a population to diverge via the following processes. Unlike a large founder population, immigration not only heightens the genetic variation that allows selection to act (Smadja & Butlin, 2011) but also increases the genetic incompatibilities that enable divergence to proceed (Seehausen, 2013) or triggers hybrid speciation (Lamichhaney et al., 2018). Moreover, if colonization occurred during a glacial period when a sea barrier was much narrower, the widened sea barrier during the subsequent interglacial period facilitates the acceleration of speciation (Carine et al., 2004; Weigelt, Steinbauer, Cabral, & Kreft, 2016). Therefore, recurrent immigration could be a key process even for the evolution of remote island endemics. A future study should include a detailed reconstruction of temporal demographic changes and phenotypic and genomic evolution in the presence of gene flow, which could be achieved via substantial sampling and the application of next-generation sequencing techniques.

### Factors affecting the specificity of colonization demography

Why were different scenarios with different influences of founder effects and recurrent immigration were supported on different islands? Isolation and island area are the two major determinants of remote island biodiversity and therefore population dynamics (MacArthur & Wilson, 1967; Valente et al., 2020). In the present study, the isolation level (the closest distance to the mainland is ~300 km between KKJ and Kyushu, ~570 km between MDT and Kyushu, and ~900 km between CCJ and Kanto) and land area (57, 30, and 24 km^2^ for KKJ, MDT, and CCJ, respectively) differed markedly. However, it remains unclear why KKJ, the closest and largest island, was colonized last, whereas the highly remote MDT was the first island colonized and, surprisingly, subject to multiple recurrent immigration events. Ecological differences are not likely the cause because pronounced differences do not exist among the three islands compared with those among the mainland and islands, including the species richness of the terrestrial breeding avifauna (15 for KKJ [Hamao & Torikai, 2011]; 9 for MDT [Takehara et al., 1999]; and 6 for CCJ [Kawakami, 2019], excluding the bull-headed shrike) and climate (humid subtropical for KKJ, tropical rainforest for MDT, and tropical monsoon for CCJ). Indeed, the ecology is similar among the islands, especially in relation to the shrikes, given that their successful colonization is possibly linked to the expansion of the anthropogenic landscape on these islands (Chiba, 1990; Matsui & Takagi, 2017).

The directionality of storms and prevailing winds can affect the frequency of immigration (Gillespie et al., 2012), potentially affecting the likelihood of recurrent immigration. Tropical cyclones (typhoons) usually pass Japan on a track from the southwest toward the northeast during autumn when seasonal migration and postfledgling dispersal occur (Japan Meteorological Agency, 2020), resulting in north winds in the western part of a cyclone that could potentially carry birds away from their normal range to southern remote islands. Indeed, an individual bull-headed shrike flying over the Pacific Ocean was sighted 500 km south of the Japanese mainland at N29° E135° after an autumn typhoon (Itakura, 1985; Figure 1a). Because the annual occurrence rate of typhoons is high around MDT, moderate around KKJ, and low around CCJ (Makino, 1986), the high frequency of immigration onto MDT may reflect the likelihood of winds carrying shrikes to the island.

The number of immigrants that affect the efficacy of founder effects may be related to seasonal migration. Our STRUCTURE analysis results suggested that the genetic cluster (cluster A) reflecting the rare allelic combination of the CCJ population was found at a relatively high level in shrikes from Hokkaido and Nagano, i.e., those located at high latitudes or high altitudes where migratory shrikes are abundant (Brazil, 2009; Endo & Ueda, 2016; Yosef & International Shrike Working Group, 2020). At low latitudinal or altitudinal regions, including Kanto, Osaka, and Kyushu, shrikes had small portions of this cluster, which is in accordance with the co-occurrence of resident and migratory shrikes (Imanishi, 2005; Yosef & International Shrike Working Group, 2020). Bull-headed shrikes are solo nocturnal migrants (S. Hara pars. obs.; Figure S6). Given the observation of a shrike on the Pacific Ocean after a typhoon by Itakura (1985), there may be many independent incidents of such solo migrating individuals being displaced far out into the Pacific Ocean by typhoons, some of which may reach remote islands and form a small founder population, as was the case on CCJ. Conversely, immigrants that caused the population establishment on KKJ and the abrupt change in the genetic structure on MDT are possibly associated with different processes such as dispersal movements by postfledging flocks consisting of several dozen juveniles (Kurata, 1967; Yamagishi, 1981). A rare migratory immigrant likely arrived at MDT, however, as reflected by one individual with 38% of cluster A (Figure 4a, arrow). The likelihood that an island receives migratory and nonmigratory immigrants as well as the proportion of such immigrants may be dependent on the migratory and dispersal routes of birds (Lees & Gilroy, 2014; Paradis et al., 1998). Our shrike study system has the potential to be used for assessing how seasonal migration contributes to the population establishment and diversification of birds (Rolland, Jiguet, Jønsson, & Condamine, Morlon, 2014).

### Conclusion

Recently established populations of animals founded by natural colonization may reflect the ecologically realistic genetic structure of the early-stage of population colonization (Clegg et al., 2002; Grant et al., 2001). In the present study, a rare system was evaluated in which birds colonized multiple remote islands 200–600 km from their normal breeding areas within only several decades. Our genetic analyses indicated that three remote islands were colonized with different demographic backgrounds and allowed us to conclude that recurrent immigration from the mainland is important for population persistence, even on remote islands. This finding is unexpected because remote island endemics are often assumed to have evolved through a rare long-distance dispersal event; however, we argue that studying the influence of gene flow at the initial stage of population divergence is crucial. To the best of our knowledge, this is the first study in which multiple recent colonization events were compared genetically at the population level. Moreover, our study bridges the gap between population genetics and macro-scale island biogeography by providing new insights into the process of population establishment.

## Data availability statement

Scripts and data for analysis are available on Dryad: https://datadryad.org/stash/share/XtfJyiIF5kFaxf7cOi6fJ12XuEct8i1mMHMpJA_9mBg. Processed genetic data (product lengths of the microsatellite alleles in the genetic dataset converted from the raw data from fragment analysis) was submitted to the data repository. Settings and procedures for these processes are fully described on the main and supplementary texts.

## Acknowledgements

Tissues and blood samples were kindly provided by Yamashina Institute for Ornithology and National Museum of Nature and Science. We thank the many members of our field teams (Masako Ueda, Sadao Imanishi, Isato Chiba, Yasui Takaya, Yuko Tsuchiya, Mariko Senda, Akari Aakamatsu, Hiroaki Matsumiya, and Seiichi Hara) and Institute for Boninology for their helps with data collection. S. Hara also kindly provided us his valuable data and information on migration of bull-headed shrikes for our supplementary material. Ayumi Matsuo and Yoshihisa Suyama gave us many useful tips for the PCR settings and genotyping protocols. We are thankful for experimental assistance from Mariko Senda. Akira Sawada, Yusuke Nishida, and Yusaku Okubo provided many useful comments on statistical analyses. The manuscript was greatly improved by an encouragement message and valuable comments by Ituro Koizumi. DNA samples were collected with permission from the Ministry of the Environment Government of Japan. This study was partly conducted with the support of a grant-in-aid for Scientific Research (C) to MT (no. 97J08530) and to DA (no. 19J21406) from the Japanese Society for the Promotion of Science (JSPS) and the Support Program of FFPRI for Researchers Having Family Obligations.

## Biosketch

Daisuke Aoki is an evolutionary ecologist, who is specialized in the fields of phylogeography, population genetics and avian migration. His research aim is to bridge a gap between microevoltuion and macroevolution by asking how biogeographic histories of organisms interacted with natural selection, stochastic processes, and evolutionary constraints. He achieves this aim through multifaceted approaches, including genetics and genomics, spatial modelling, mathematical simulation, and field biologging approaches.

DA, SM and MT conceived the ideas and designed the study; DA, SM, MT, and IN led the fieldwork; DA led the laboratory procedures with supports by ME, JN, and IN; DA analysed the data, coordinated the study, prepared the draft, led the writing and revised the manuscript with assistance from MT and SM. All authors gave final approval for publication and agree to be held accountable for the work performed therein.

