## Supporting Materials for "Population genetics of recent colonization suggests the importance of recurrent immigration on remote islands"

Affiliations

**Supplementary methods**

*Experimental and genotyping protocols*

We firstly dried up 1 μL of the template DNA solution (containing approximately 20 ng of DNA) in the reaction well, by using a thermal cycler, and polymerase chain reactions (PCRs) were performed by directly adding reaction mixtures prepared in the well with DNA dried up at its bottom. Reaction mixtures were separately prepared for the three different multiplex sets (see Table S2) in total volume of 2 μL, containing 2× Multiplex PCR Master Mix (Qiagen) and primer mixtures containing primer sets prepared for one of the multiplex sets. Primer sets were consisted of 0.2 μM of each forward primer (containing primers with and without fluorescence labels in a specific ratio) and 0.2 μM of each reverse primer that was PIG-tailed (Brownstein et al., 1996). The ratio between the fluorescent and non-fluorescent forward primers were individually determined for each set of primers (see Table S2). The PCR started with an initial denaturation step at 95 °C (15 min), 38 cycles of series of denaturation at 94 °C (30 s), annealing at specific annealing temperature for each primer (1.5 min), and extension at 72 °C (1.0 min) and a final extension step at 72 °C (30 min). PCRs were all performed using the Veriti 200 thermal cycler (Applied Biosystems). PCR products were analysed using ABI 3730 Genetic Analyzer (Applied Biosystems) with GeneScan 600 LIZ dye Size Standard v 2.0 (Applied Biosystems). Peak Scanner software v 1.0 (Applied Biosystems) was used to analyse the lengths of fragments. If loci were not amplified or fragment lengths were ambiguous, they were subjected to repeated PCRs and fragment analyses. Repeated PCRs were conducted with total volume of 5 μL of the reaction mixtures.

*Sibship assignment by COLONY*

We inferred pairs of individuals in relationships of full-siblings amongst samples collected at the same local population by COLONY v.2.0.6.6 (Jones & Wang, 2010), and we retained one of the individuals assigned to each pair of the putative full-siblings. We assumed monogamy for both males and females while priors on parents were not defined. Full-siblings were determined when probabilities of sib-ship exceeded 95%. If multiple of individuals were assigned to a group of full-siblings, only one individual was randomly selected to be retained per each group in the dataset. There is a case wherein one female parent shrike and its offspring (which was supported by field observation) were grouped together as a set of “full-siblings” in the analysis. This does not cause any problem in our genetic analyses because data without relatedness was generated to avoid the genetic structure reflecting specific kin-structure, which violates some assumptions of the downstream analyses. In case that both adults and yearlings were included in the sibship groups, adult samples were prioritized to be retained in the data without relatedness.

*Genetic structure predictions on the temporal samples for MDT*

For the STRUCTURE analysis, we used the option “PFROMPOPFLAGONLY.” When the MDT samples collected in 2002–2008 and the remaining samples used in the previous analysis were assigned with Popflag = 0 and Popflag = 1, respectively, the results of STRUCTURE analysis with and without the additional samples were similar but the genetic structure of the additional samples was simultaneously predicted. *K* = 4 was used, the option “USEPOPINFO” was turned off, and the alpha value was fixed after it was set as the mean of 10 runs of the previous analyses (alpha = 0.066). For the DAPC, additional samples were predicted by using the previous results described in the main texts.

**Supplementary tables**

**Table S1.** Information on sampling localities and dates

| **Region** | **N** | **Locality (Coordinates)** | **Collection periods** | **Collection Number  *tissues at YIO or NSMT** | **Source** |
| --- | --- | --- | --- | --- | --- |
| Hokkaido | 32 | Ishikari City, Hokkaido Pref. (43.19, 141.36) | Jul. 1997,  May 1998 ~ Jun. 1998 | - | MT |
| Nagano | 25 | Matsukawa Town, Nagano Pref.  (35.60, 137.90) | May 2019 ~ Jun. 2019 | - | DA |
| Osaka | 54 | Sakai City, Osaka Pref.  (34.57, 135.53) | Mar. 1989 ~ Jun. 1989 | NSMT1 – NSMT166 (Ad or 1Y) | NSMT (IN) |
| Chichijima | 18 | Ogasawara Vlg., Tokyo Pref.  (27.08, 142.22) | May. 1995  Apr. 1997 ~ Jun. 1997  Feb. 1998 ~ Mar. 1998 | 1995-0643 | MT, YIO |
| Kikaijima | 5 | Kikai Town, Kagoshima Pref.  (28.33, 129.97) | Jul. 2012  Apr. 2017 ~ May 2017 | NSMT51578  NSMT54371  NSMT54372  NSMT54373  NSMT54374 | NSMT |
| Kita-Daito Island | 4 | Kita-Daito Vlg.,  Okinawa Pref.  (25.94, 131.31) | May 2008 ~ Jul. 2008 | - | SM |
| Minami-Daito Island (1998) | 30 | Minami-Daito Vlg., Okinawa Pref.  (25.84, 131.24) | Jan. 1998 ~ May 1998 | - | MT |
| Minami-Daito Island (2002) | 58 |  | Apr. 2002 ~ Nov. 2002 | - | SM, MT |
| Minami-Daito Island (2003) | 80 |  | Jan. 2003 ~ Nov. 2003 | - | SM |
| Minami-Daito Island (2004) | 43 |  | Mar. 2004 ~ Aug. 2004 | - | SM |
| Minami-Daito Island (2005) | 59 |  | Mar. 2005 ~ Nov. 2005 | - | SM |
| Minami-Daito Island (2006) | 38 |  | Feb. 2006 ~ Jul. 2006 | - | SM |
| Minami-Daito Island (2007) | 43 |  | Feb. 2007 ~ Jul. 2007 | - | SM |
| Minami-Daito Island (2008) | 44 |  | Feb. 2008 ~ Jul. 2008 | - | SM |
| Kanto | 18 | 36.2, 139.7  36.2, 139.7  35.7, 139.9  35.9, 140.1  36.0, 139.9  35.9, 140.1  35.9, 140.1  36.0, 139.4  36.1, 140.1  36.0, 139.9  36.0, 139.9  36.0, 139.9  35.7, 139.7  35.4, 138.9  35.5, 139.3  35.5, 138.8  35.4, 139.4  35.4, 139.3 | Dec. 2004  Sept. 2004  Mar. 1997  Jun. 2012  Oct. 2012  May 2006  May 2006  Dec. 1997  Feb. 2001  Nov. 2005  Nov. 2005  Nov. 2005  Oct. 2015  Oct. 2009  Jun. 2009  Jan. 2012  Oct. 2010  Oct. 2010 | 2004-5027  2004-5266  1997-0081  2012-0222  2012-1844  2012-2308  2012-2309  1997-0482  2001-0056  2017-0439  2017-0456  2017-0485  2015-0900  2011-0705 | YIO, DA |
| Kyushu | 14 | 34.5, 129.3  33.7, 129.7  32.1, 130.3  32.1, 130.3  32.1, 130.3  32.1, 130.3  32.1, 130.3  32.1, 130.3  31.8, 130.3  31.8, 130.3  32.4, 131.5  32.4, 131.5  32.1, 131.5  31.8, 131.4 | Nov. 2007  Feb. 2004  Feb. 1999  Feb. 2017  Feb. 2017  Feb. 2010  Feb. 2010  Feb. 2010  Feb. 2011  Feb. 2011  Feb. 2007  Feb. 2007  Feb. 2007  Jan. 2007 | 2008-0052  2004-5062  2000-0041  2017-0367  2017-0368  2010-0193  2010-0198  2010-0269  2011-0404  2011-0430  2007-0678  2007-0695  2007-0296  2007-0500 | YIO |

**Table S2.** Primer sequences, primer combinations and annealing temperatures in each multiplex PCR set. The sequence shaded by grey corresponds to an additional part of the PIG-tail motif added to the original sequence. The forward and reverse primers defined here do not correspond to the original descriptions since decision on primers to be labelled were uniquely made

| **Locus** | **Primer sequences** | **Dye** | **ref** | **Ratio** [fluorescent:  non-fluorescent] | **Multiplex Set** | **Annealing temperature [°C]** |
| --- | --- | --- | --- | --- | --- | --- |
| SJR4 | 5’-TCCAGGCTGTGCTTGCACTTG-3’ / 5’-GTTTCTTGCCAGACCACCAACTAAATC-3’ | PET® | Mundy et al. (1997) | 1:24 | 1 | 57.5 |
| CoBr02 | 5’-AGGGAAACGGCTGGAAAACTTG-3’ / 5’-GTTTCTTGCCCTCATCGATAGCGTACATATC-3’ | NED® | Schoenle et al. (2006) | 1:30.7 | 1 | 57.5 |
| Llu55 | 5’-AGTTGGGCACCACAGAAAGC-3’ / 5’-GTTTCTTCCTGATGGGTCCTTCCAAC -3’ | PET® | Coxon et al. (2012) | 1:16 | 1 | 57.5 |
| CoBr09 | 5’-AGGCCGATTTCTTGCTTTTC-3’ / 5’-GTTTCTTCAATGCCATCAGGACAAGTAAAC-3’ | 6-FAM® | Schoenle et al. (2006) | 1:3.35 | 1 | 57.5 |
| LOX1 | 5’-CCACACACATTCACTCTATTG-3’ / 5’-GTTTCTTATGATGGTAAGTCTAATGAAAGC-3’ | PET® | Piertney et al. (1998) | 1:1.45 | 1 | 57.5 |
| Llu82 | 5’-GGCTATAAATCTGATGTAACCC-3’ / 5’-GTTTCTTGCTTAGTGCATTATGTGTTTCC-3’ | VIC® | Coxon et al. (2012) | 1:33 | 2 | 58.1 |
| LTMR7 | 5’-CCCACCACCCTCCACCTTGTT-3’ / 5’-GTTTCTTACTGTCACCTGTTGCTTTCCA-3’ | NED® | Bhagabati et al. (2004) | 1:22 | 2 | 58.1 |
| ApCo15 | 5’-TGTGCCTCAGTATACCATCAGT-3’ / 5’-GTTTCTTTTCAGAGTCTCTGTCTTAGCTG-3’ | 6-FAM® | Stenzler and Fitzpatrick (2002) | 1:24 | 2 | 58.1 |
| LS1 | 5’-AACTCTTTTGCTCTCTCCCTATACC-3’ / 5’-GTTTCTTCTCTTGATTCATTCAGTGGACACC-3’ | PET® | Mundy et al. (1997) | 1:12 | 2 | 58.1 |
| Llu102 | 5’-TAAGACTTGGTGGGTTATTGC-3’ / 5’-GTTTCTTCCCAACCTTGCAAATTCA-3’ | NED® | Coxon et al. (2012) | 1:31 | 2 | 58.1 |
| Llu176 | 5’-CTGAACTCACTAAAGTCTGC-3’ / 5’-GTTTCTTGAATTAGGCAAGCAGTTGG-3’ | VIC® | Coxon et al. (2012) | 1:64.7 | 3 | 57.5 |
| LS2 | 5’-CTCAAACTGAAAATGTAGATTCACC-3’ / 5’-GTTTCTTTACATTTTTTCCCATGAGGC-3’ | PET® | Mundy et al. (1997) | 1:26 | 3 | 57.5 |
| Llu39 | 5’-CAGCCCTGAATTGGTCTGGC-3’ / 5’-GTTTCTTCCACCTTCAGTTGCAGTTTCC-3’ | 6-FAM® | Coxon et al. (2012) | 1:18.5 | 3 | 57.5 |
| Llu95 | 5’-GATGAGTGTGAGGGAATGAC-3’ / 5’-GTTTCTTCACATAAATCCCAAGTCCTC-3’ | NED® | Coxon et al. (2012) | 1:17.9 | 3 | 57.5 |
| CoBr25 | 5’-CCAGGAGGCAAACAGCACTTAC-3’ / 5’-GTTTCTTCCCAGAAGGCTCCAACATTATC -3’ | VIC® | Schoenle et al. (2006) | 1:28 | 3 | 57.5 |

**Table S3.** Null allele frequency estimates calculated for each locus and each population. Values higher than 0.2 are shown in bold italic. NA indicates that null allele frequency could not be estimated. The dataset without relatedness was used.

|  | **Hokkaido** | **Kanto** | **Nagano** | **Osaka** | **Kyushu** | **CCJ** | **KKJ** | **MDT 1998** | **MDT 2002** | **MDT 2003** | **MDT 2004** | **MDT 2005** | **MDT 2006** | **MDT 2007** | **MDT 2008** | **KDT** |
| --- | --- | --- | --- | --- | --- | --- | --- | --- | --- | --- | --- | --- | --- | --- | --- | --- |
| **SJR4** | 0.000 | 0.000 | 0.068 | 0.000 | 0.085 | 0.000 | 0.000 | 0.000 | 0.019 | 0.004 | 0.000 | 0.000 | 0.000 | 0.000 | 0.000 | ***0.202*** |
| **CoBr02** | 0.000 | 0.000 | 0.000 | 0.000 | 0.134 | 0.001 | 0.001 | 0.029 | 0.000 | 0.000 | 0.053 | 0.000 | 0.000 | 0.142 | 0.000 | 0.000 |
| **Llu55** | 0.025 | 0.116 | 0.030 | 0.086 | 0.066 | 0.066 | 0.000 | ***0.225*** | ***0.230*** | ***0.263*** | ***0.256*** | ***0.250*** | 0.173 | ***0.254*** | ***0.247*** | ***0.387*** |
| **CoBr09** | 0.000 | 0.000 | 0.040 | 0.000 | 0.027 | 0.000 | 0.000 | 0.000 | 0.059 | 0.010 | 0.000 | 0.000 | 0.080 | 0.019 | 0.056 | 0.000 |
| **LOX1** | 0.000 | 0.000 | 0.000 | 0.017 | 0.000 | 0.000 | 0.000 | 0.000 | 0.000 | 0.000 | 0.000 | 0.000 | 0.000 | 0.000 | 0.000 | 0.001 |
| **Llu82** | 0.037 | 0.000 | 0.036 | 0.011 | 0.000 | 0.000 | 0.157 | 0.017 | 0.000 | 0.000 | 0.016 | 0.000 | 0.000 | 0.000 | 0.000 | 0.000 |
| **LTMR** | 0.000 | 0.009 | 0.000 | 0.018 | 0.000 | 0.000 | 0.000 | 0.010 | 0.000 | 0.000 | 0.009 | 0.036 | 0.000 | 0.000 | 0.000 | 0.000 |
| **ApCo15** | 0.030 | 0.030 | 0.000 | 0.000 | 0.073 | 0.000 | 0.016 | 0.015 | 0.002 | 0.000 | 0.027 | 0.001 | 0.000 | 0.038 | 0.000 | 0.000 |
| **LS1** | 0.000 | 0.000 | 0.000 | 0.028 | 0.000 | 0.000 | 0.072 | 0.000 | 0.045 | 0.040 | 0.000 | 0.000 | 0.083 | 0.014 | 0.000 | 0.000 |
| **Llu102** | ***0.226*** | 0.135 | 0.176 | ***0.220*** | 0.000 | 0.097 | ***0.269*** | 0.104 | ***0.220*** | ***0.223*** | 0.175 | 0.197 | ***0.234*** | ***0.203*** | 0.162 | 0.132 |
| **Llu176** | 0.000 | 0.000 | 0.102 | 0.070 | 0.000 | 0.078 | 0.000 | 0.000 | 0.006 | 0.128 | 0.179 | 0.072 | 0.106 | 0.155 | 0.040 | 0.083 |
| **LS2** | 0.000 | 0.000 | 0.007 | 0.026 | 0.063 | 0.000 | 0.200 | 0.000 | 0.000 | 0.000 | 0.001 | 0.000 | 0.000 | 0.000 | 0.000 | 0.001 |
| **Llu39** | 0.026 | 0.057 | 0.000 | 0.011 | 0.000 | 0.046 | 0.000 | 0.000 | 0.000 | 0.035 | 0.000 | 0.000 | 0.000 | 0.062 | 0.028 | 0.000 |
| **Llu95** | 0.032 | 0.039 | 0.041 | 0.000 | 0.000 | 0.000 | 0.000 | 0.000 | 0.017 | 0.026 | 0.000 | 0.023 | 0.000 | 0.008 | 0.000 | 0.000 |
| **CoBr25** | 0.000 | 0.061 | 0.071 | 0.000 | 0.073 | 0.076 | 0.000 | 0.000 | 0.000 | 0.000 | 0.057 | 0.055 | 0.000 | 0.000 | 0.000 | 0.000 |

**Supplementary figures**

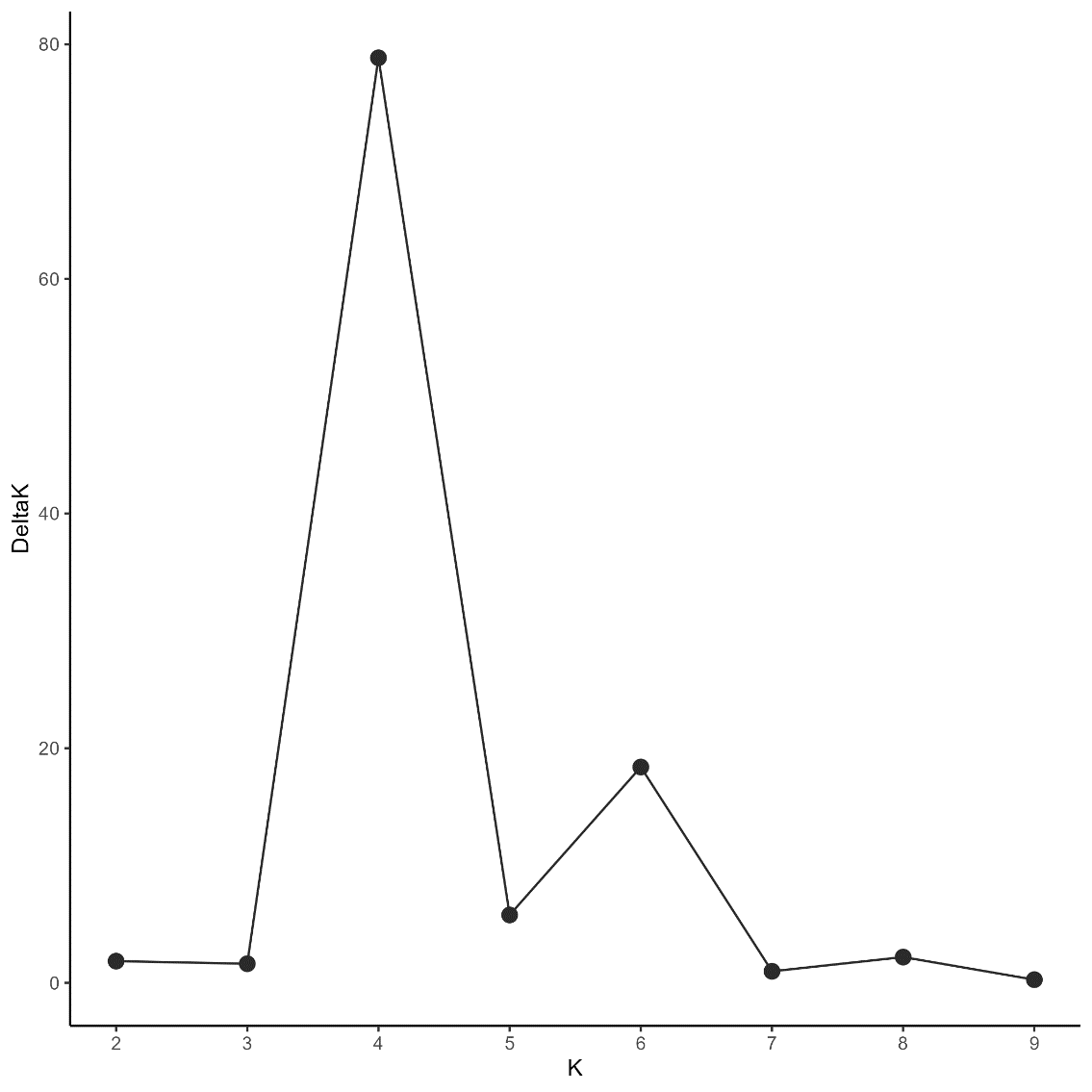

**Figure S1.** Delta K calculated for each K by the Evanno method.

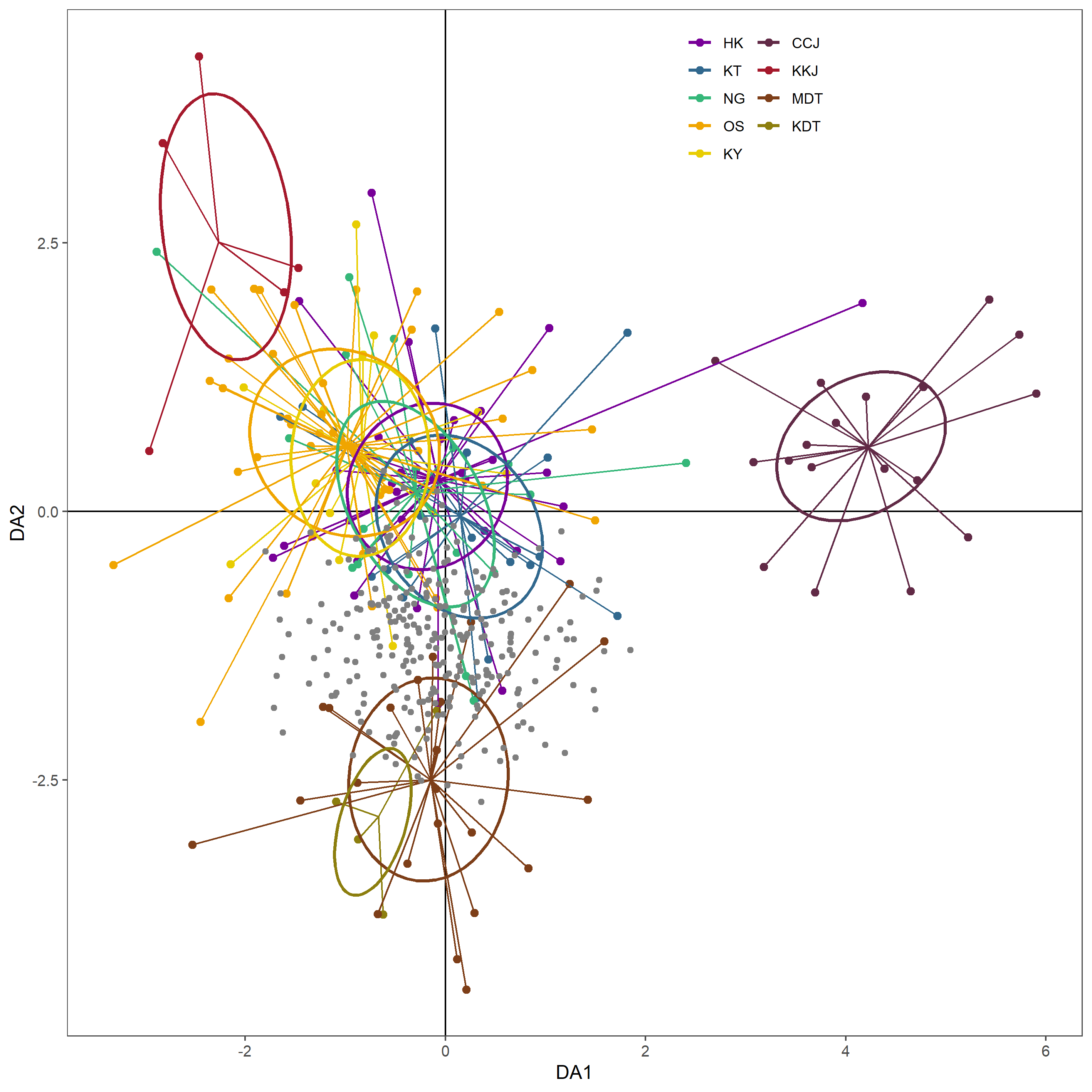

**Figure S2**. Positions of the Minami-Daito Island (MDT) samples collected between 2002 and 2008 (grey dots) on the DAPC plane (same as figure 4b).

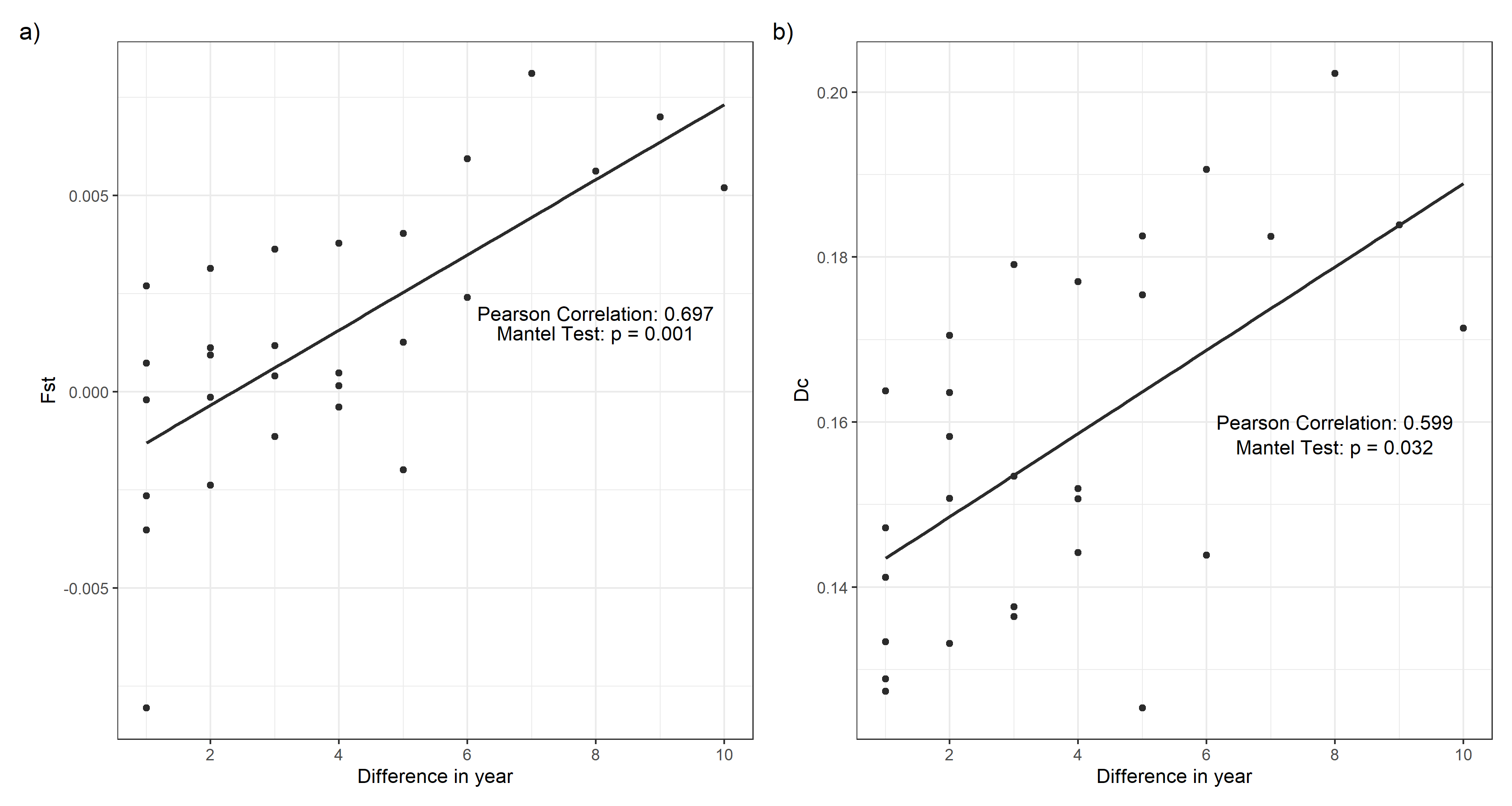

**Figure S3.** Temporal change in the genetic structure of bull-headed Shrikes collected from 1998 to 2008 on Minami-Daito Island. (a) Pairwise Fst and (b) Cavalli-Sforza and Edwards’ genetic distance (Dc) are used as the y-axis, and the x-axis shows pairwise differences in years between samples.

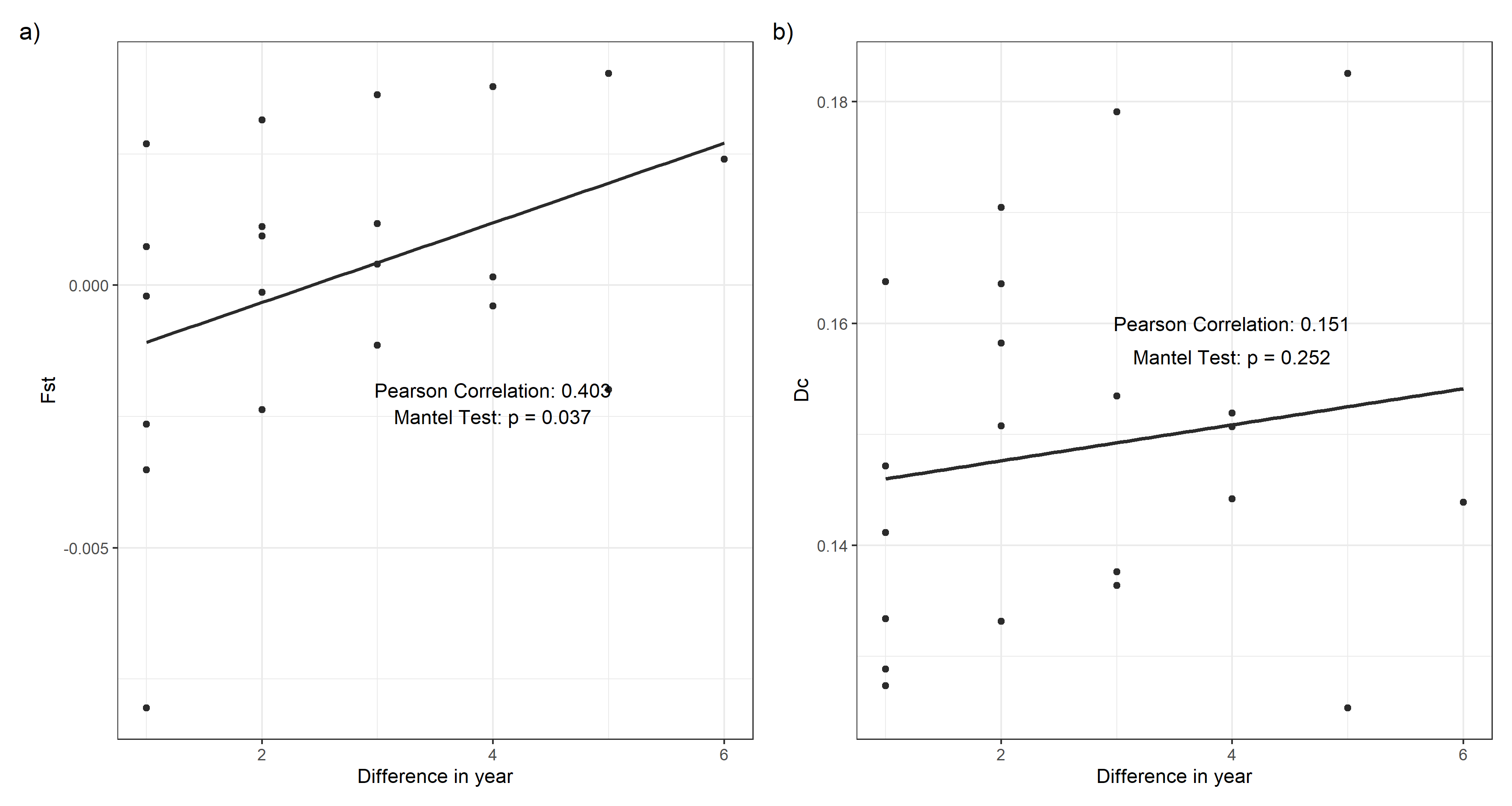

**Figure S4.** Temporal change in the genetic structure of bull-headed shrikes collected from 2002 to 2008 on Minami-Daito Island, with excluding samples collected in 1998. (a) Pairwise Fst and (b) Cavalli-Sforza and Edwards’ genetic distance (Dc) are used as the y-axis, and the x-axis shows pairwise differences in years between samples.

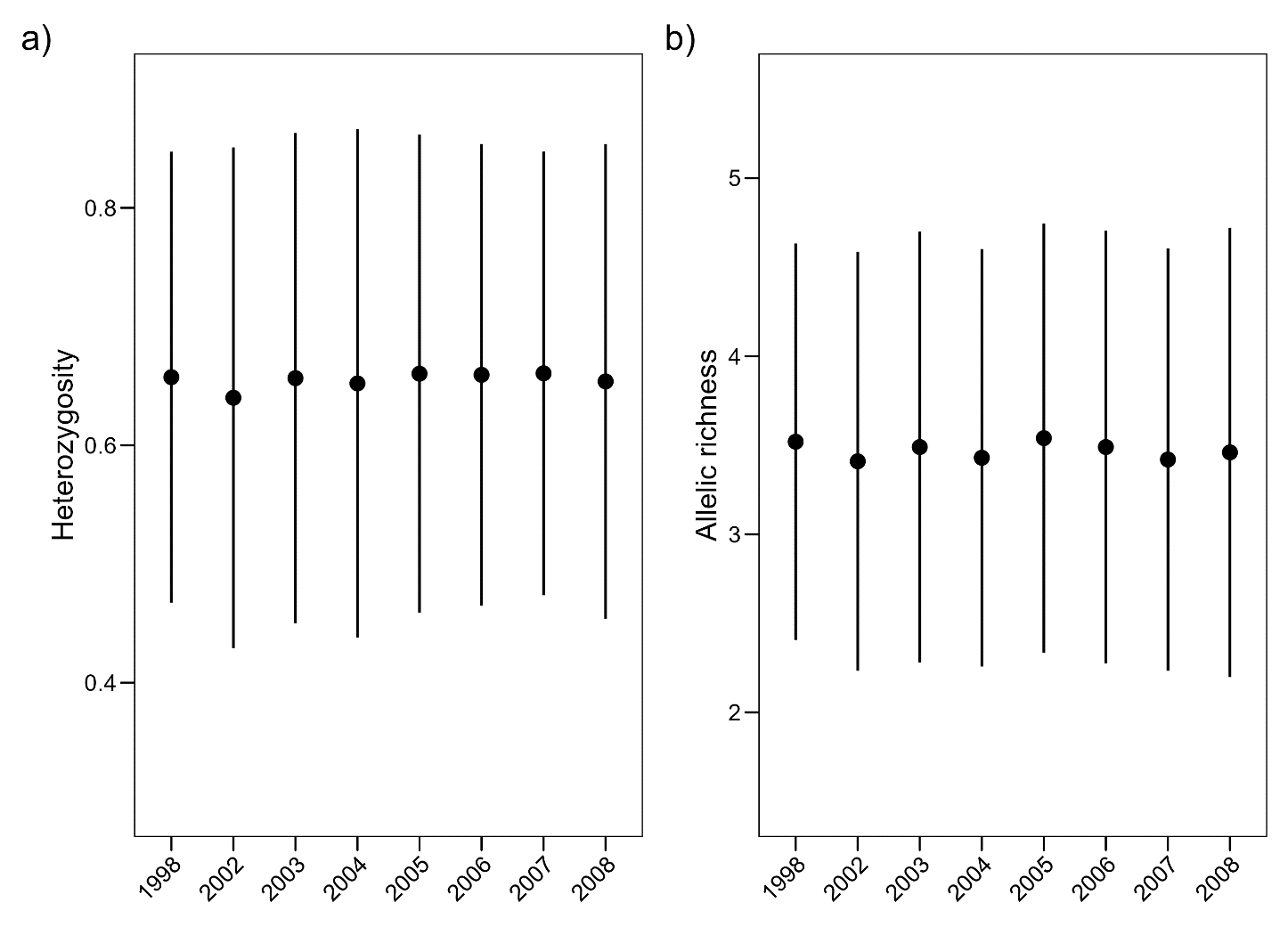

**Figure S5.** Heterozygosity and allelic richness of the samples collected on Minami-Daito Island in different years

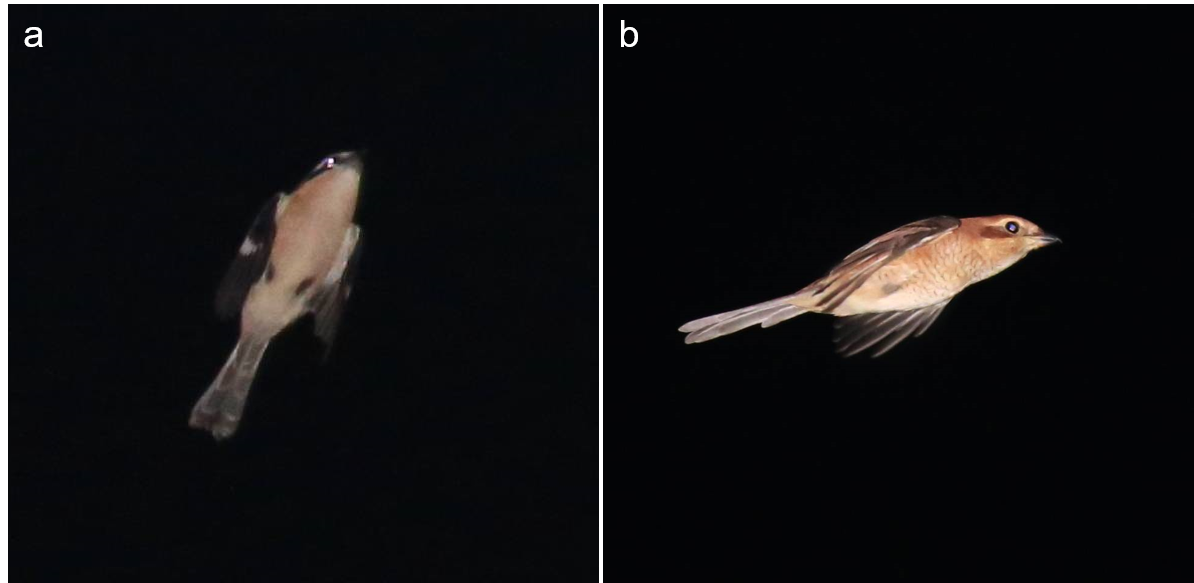

**Figure S6** Photos of nocturnal and solitary flight of migration of a a) male and b) female bull-headed shrikes. Photos were kindly provided by and used with permission of Seiichi Hara
